# In situ light-driven pH modulation for NMR studies

**DOI:** 10.1101/2025.01.16.633412

**Authors:** Aarav Barde, Ruixian Han, Martin A. Olson, Marco Tonelli, Chad M. Rienstra, Katherine Henzler-Wildman, Thirupathi Ravula

## Abstract

Proton exchange is a fundamental chemical event, and NMR provides the most direct readout of protonation events with site-specific resolution. Conventional approaches require manual titration of sample pH to collect a series of NMR spectra at different pH values. This requires extensive sample handling and often results in significant sample loss, leading to reduced signal or the need to prepare additional samples. Here, we introduce a novel approach to control pH in NMR samples using water soluble photoacids, which alter the pH of the solution from near neutral to acidic pH upon in situ photo-illumination. We show that the solution pH can be precisely controlled by choice of illumination wavelength and intensity and sufficient protons are released from the photoacid to achieve meaningful pH change in samples where the molecule of interest has significant buffering capacity, such as a >100 μM protein sample. The pH is monitored *in situ* using internal standards with pH-sensitive chemical shifts. This method enables precise, calibrated, non-invasive change of sample pH within an NMR magnet, dramatically reducing the necessary sample handling. These findings highlight the potential of light-induced pH control in NMR experiments and increase the robustness and reliability of pH-dependent studies. With pH playing a key role in modulating chemical behavior in both biological and synthetic systems, the ability to study protonation states and modulate sample pH in a simple and precise manner that is compatible with high-resolution NMR studies of molecular structure and function has wide applications.

**Entry for the Table of Contents:** In this work, we introduce a novel approach to control pH in NMR samples using light-activated photoacids. By combining light stimuli with pH-sensitive molecules, we demonstrate the ability to precisely modulate pH without physically manipulating the sample. This method enables non-invasive pH titration in NMR studies, where pH plays a key role in protein function. Our findings highlight the potential of light-induced pH control to overcome existing limitations in NMR, providing a powerful tool for advancing protein research under controlled pH conditions.

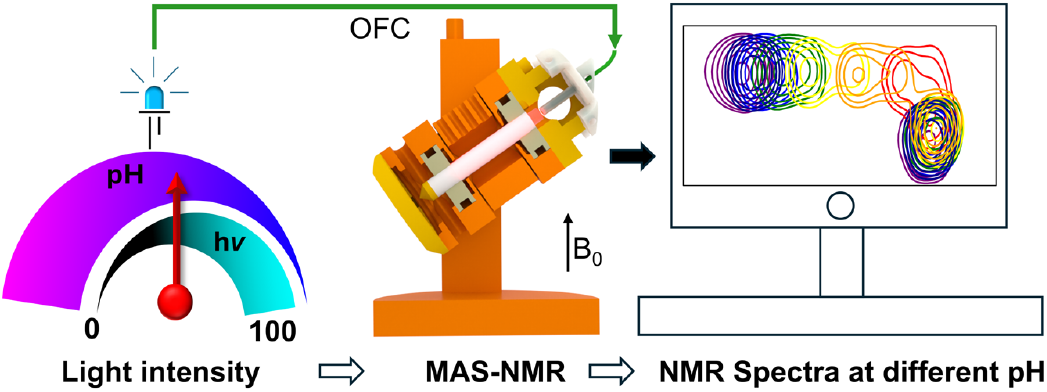

---

NMR is a versatile technique for studying the structure, dynamics, and chemistry of molecules ranging from small molecules to biological macromolecules. pH is an important variable affecting the protonation state of functional groups that impacts the structure and reactivity of these molecules. NMR is one of the few methods that can directly detect protonation events through the impact of protonation on the chemical shift of the protonatable group and surrounding nuclei.^[1-3]^ However, controlling and measuring pH in small volumes within NMR samples is challenging, particularly for small-volume solution NMR samples in narrow tubes or for solid-state NMR samples packed into rotors with microliter volumes requiring hermetic sealing.^[2]^ Significant sample losses can occur if the sample is repeatedly removed from the tube or rotor to change the pH or multiple samples must be prepared for titration. However, many interesting biological molecules cannot be produced in the large quantities needed for this approach,^[4]^ creating a major bottleneck for such studies. Other methods, such as pKa determination using non-aqueous solvents, have been shown to successfully measure the pKa of small organic molecules within a single sample,^[5]^ however no such methods currently exist for biological systems under aqueous conditions.

Using light stimuli as a non-invasive technique to control pH would be a significant breakthrough, enabling pH manipulation and titration of proteins or other molecules in NMR rotors or tubes without the need to unpack and repack samples or prepare multiple samples. This can be achieved using photoacids^[6]^, small molecules that bind or release protons upon photo-illumination. Light-activation of caged compounds and light-based control of optogenetic tools have become widespread and demonstrate the power of using light to control biochemical processes, regulate pH or small molecule concentrations and rapidly tune environmental conditions within cells^[7]^.Availability of different photoacids further extends the accessible pH range to span values relevant for activation of membrane proteins,^[8]^ DNA assembly^[9, 10]^ and other biological applications.^[6, 11]^ The simple and precise control of light intensity and wavelength underlies its power and versatility to manipulate and study conditions with high spatial and temporal resolution in locations that are difficult to access directly.^[12,13]^ These tools are readily adaptable to other locations where direct manipulation is challenging, such as samples in an NMR spectrometer.^[14]^ It has already been shown that photoacid induced pH changes can be achieved by in-situ illumination under solution NMR conditions with high accuracy using ultraviolet light. However, this method has not been applied to biological samples due to the limitation of strong Xenon lamp radiation that can damage protein samples and the irreversible nature of the photoacid used in these studies.^[15]^

Biological macromolecules, such as proteins, biological extracts, and complex reaction mixtures will frequently have multiple protonatable groups and relatively high μM-mM concentrations are generally required for NMR studies. Thus, the sample itself often has significant buffering capacity. Prior demonstration of light-induced pH change using a photoacid only included the NMR pH sensor molecule (histidine) and photoacid in the NMR tube. For this technology to be broadly useful, highly soluble photoacids are required that can release sufficient protons to achieve meaningful pH changes in the presence of the highly self-buffered sample of interest. Here we present a new photoacid that meets these requirements to achieve pH control under strong buffering conditions and achieve light-controlled pH change of ≈2 pH units in protein NMR samples.

We combine in-situ photo illumination of the photoacid with pH sensor molecules that allow us to read out the pH through chemical shift changes to achieve accurate pH titration in solid-state NMR with a single sample. We used Immunoglobulin-binding domain B1 of streptococcal protein (GB1) as an example to demonstrate reversible, light-controlled pH titration within the solid-state NMR rotor, since this provides a more challenging system for pH titration than solution NMR^[16]^ due to the difficulty in unpacking and repacking solid-state rotors to change sample pH, the higher cost (both rotor cost and greater sample requirement) to prepare multiple samples for a solid-state NMR titration, and more challenging illumination within the spectrometer. We achieve sample illumination with minimal modification of a conventional magic angle spinning (MAS) NMR probe. Although demonstrated in a solid-state MAS rotor, our novel photoacid is readily adaptable to solution NMR experiments. Thus, these advancements open up new methods for studying the impact of pH on the structure and dynamics of biological molecules with either solid-state or solution NMR.

In-situ illumination of NMR samples under MAS is particularly challenging due to the presence of the spinning module. Two approaches have been used previously to perform in-situ illumination in MAS probes. The first method involves illumination on the side of the rotor, requiring a major redesign and expensive transparent rotors.^[17]^ The second approach is to illuminate through the cap of the rotor, which requires minimal changes to the probe and eliminates the need for transparent rotors.^[18]^ We used the second method, employing transparent caps for light illumination. We chose to modify a Phoenix NMR HFXY 3.2 mm probe to implement this because the Phoenix probe design allows easy access to the top cap of the rotor for fiber optic illumination. We designed a 3D-printed adapter to hold the optical fiber in place, as shown in Figure 1. The rotor caps are made up of PMMA(poly(methyl methacrylate)), as they are transparent in the wavelength range needed to excite the photoacid (visible to near ultra-violet). The MAS module typically uses red light (∼650 nm) for detecting the spinning rate of the rotor. To avoid interference from sample illumination in the visible range, we substituted the red tachometer signal with much longer wavelength near infrared (NIR). The photodiode inside the NMR probe for tachometer signal detection is usually broadband and can detect NIR wavelengths, thus only the tachometer optical fiber input required modification. Overall, our design requires two optical fibers to be introduced into the magnet: one for the 980 nm NIR tachometer signal, and one for the desired frequency of sample illumination (for additional detail see supporting information figure S1).

**Figure 1.**
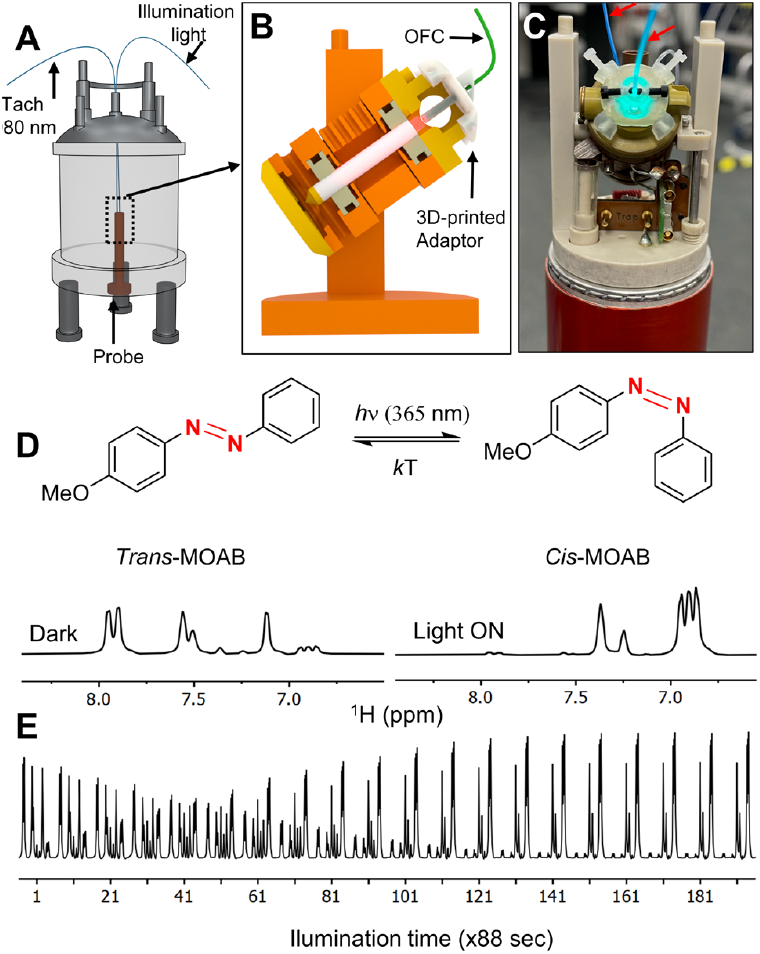
Overview of in-situ photo-illumination inside the solid-state NMR system. A) Schematic representation of an NMR magnet showing the setup of the probe and optical fiber cable (OFC) setup. B) MAS-stator model showing the 3D-printed adapter used to hold the optical fiber in place to illuminate the sample through the rotor cap using a PMMA transparent cap. C) Picture of MAS module with a 3D-printed adapter and two connected optical fibers indicated with red arrows. The sample is illuminated with 500 nm LED light. D) Chemical structure of MOAB showing its *cis*- and *trans*-conformations (top). Bottom: 1D-^1^H NMR spectra acquired under dark conditions and upon illumination with 365 nm light, demonstrating the conversion of the *trans*-conformation to the *cis*-conformation. E) Selected aromatic region of the 1D-^1^H NMR spectra acquired as a function of time upon illumination with 365 nm light, showing the conversion process over time. All the spectra were acquired on a 600 MHz instrument with MAS of 10 kHz.

First, we tested the sample illumination with the conformational conversion of a small molecule, 4-Methoxyazobenzene (MOAB). Under dark conditions, this molecule primarily exists in the trans conformation, and upon illuminating with 360 nm wavelength light, it converts into the cis conformation.^[19]^ We recorded 1H NMR spectra of a MOAB solution in deuterated Methanol (10 mg/mL) in the dark and with illumination at 365 nm. Figure 1D shows the 1H NMR spectrum as a function of time upon illumination. In the dark, the 1H NMR spectra of MOAB show a predominantly trans conformation. Upon illumination, the population of cis conformation increased until reaching complete conversion after about 4 hours. These results show that our approach fully illuminates the contents of the MAS stator.

Then, to manipulate pH within the MAS rotor, we chose to use water soluble merocyanine (MCH) photoacid, which predominantly exist in the *trans*-conformation in the absence of light. Upon illumination with 500 nm light, *trans*-MCH converts to *cis*-MCH, which then undergoes ring closure to form the spiropyran (SP) conformation, releasing a proton (Figure 2A). In the dark, the SP state of the molecule reverts to a deprotonated open structure (MC), which abstracts a proton from bulk water to returns to the original protonated MCH form. The population of these conformations is dependent on the solution pH, photoacid pKa, light and temperature.^[20]^ The pKa, solubility, and stability of the photoacid in water depends on the substitutions of the indole and phenol group moieties.^[21]^ It has been shown that the OMe group on the indole ring stabilizes the photoacid against hydrolysis,^[22]^ whereas sulphonic acid (-SO_3_ ^-^) substitution on the phenol ring increases solubility in aqueous solutions.^[8, 23]^ Since changes in pH under high-buffering conditions require sufficient photoacid solubility, we incorporated these two functional groups to enhance the solubility and stability of the photoacid as shown in Figure 2A. After synthesis, the photoacid was fully characterized using solution NMR (Figure S2 and S3), confirming its structure. The resulting apparent pKa of this photoacid was measured using UV spectroscopy and found to be 6.5 (figure S4). The apparent pKa of MCH determined from UV spectra are pseudo pKa values due to the equilibrium between the MC and SP forms, but are useful in estimating the effective concentration of the active MCH form and defining the working pH range for the photoacid.^[24]^

**Figure 2.**
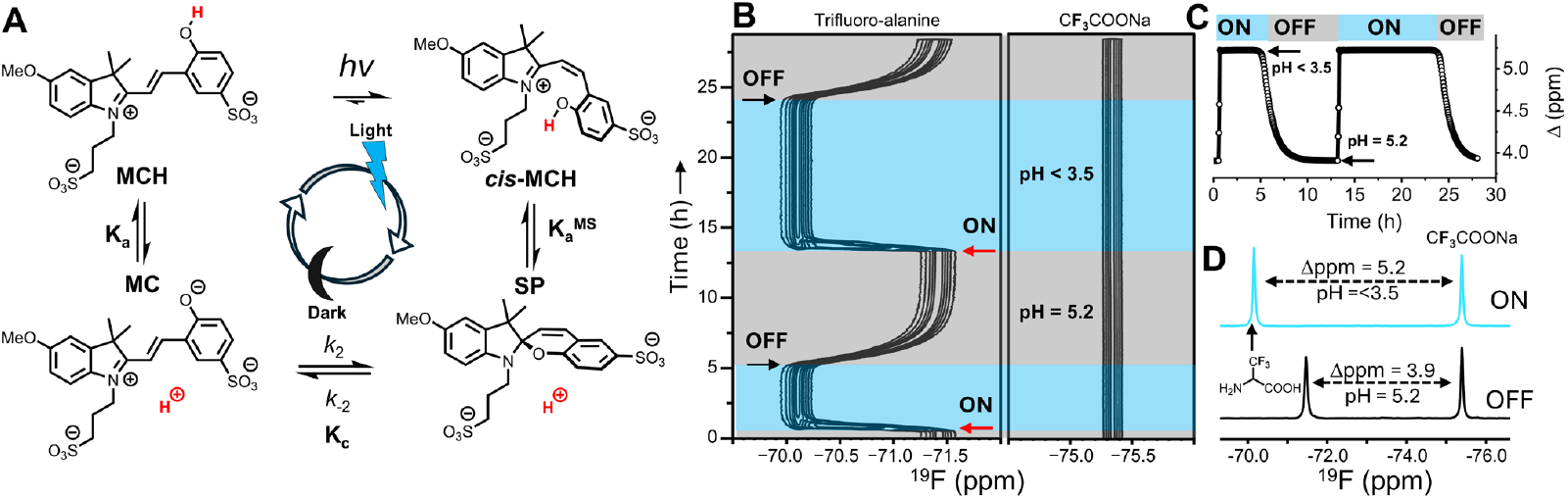
Photoacids for pH control using light: A) Chemical structure of the photoacid and its four-state model depicting proton release upon light illumination, and reversible proton uptake in the dark. B) Pseudo-2D ^19^F spectra recorded as a function of time, under dark (light off, shown in gray) and light conditions (500 nm, light on, represented in light blue). C) Plot showing the chemical shift difference between trifluoro-alanine and TFANa during light illumination as a function of time, demonstrating the pH shift from 5.2 to <3.5, and the reversibility of this process in dark conditions (light off). D) 1D-^19^F NMR spectra showing pH changes upon sample illumination with 500 nm light. Trifluoro-alanine was used as a pH sensor molecule, and sodium trifluoroacetate as a chemical shift reference. The differences in chemical shifts were used to calculate the pH of the solution. Peak assignments are shown in the figure. All the spectra were acquired with MAS of 10 kHz.

Once the appropriate photoacid was synthesized, we needed to measure the pH change achieved under in-situ illumination in the MAS rotor. To accurately measure pH inside the NMR rotor, we used trifluoro-alanine (pKa 5.6) as an NMR pH sensor and sodium trifluoro-acetate (TFANa) as the internal chemical shift reference.^[25]^ By measuring the ^19^F chemical shift difference between trifluoro-alanine and TFANa peaks, we could calculate the exact pH inside the NMR rotor (figure S5). We then prepared a sample containing 20 mM photoacid, 10 mM trifluoro-alanine, and 10 mM TFANa in a 3.2 mm rotor capped with a transparent PMMA cap. Figure 2D shows the ^19^F spectrum recorded under dark conditions, indicating a pH of 5.2, consistent with the pH measurements of the sample taken outside the rotor using a pH meter. Upon illumination with 500 nm light, the ^19^F chemical shift difference was 5.2 ppm, corresponding to a pH of less than 3.5, which is the detection limit of this pH sensing method. ^1^H NMR also showed complete conversion of MCH to SP (Figure S6). This demonstrates that we can change the pH inside the NMR spectrometer using light to release protons from our novel photoacid. Figure 2B presents pseudo-2D spectra recorded as a function of time under both light and dark conditions. The chemical shift separation plot shows that the pH dropped below 3.5 within a few minutes (time constant ∼4 min, Figure S7) of illumination and remained stable as long as the light was on. Upon turning off the light, the pH returned to its starting value (5.2) within two hours (time constant ∼ 64 min, Figure S7). This experiment was repeated, and in the second cycle, the low pH state was maintained stably for more than 10 hours with illumination (Figure 2C). These results show that light-control of pH inside the NMR rotor can be stably maintained over the timescales needed to perform multidimensional NMR experiments.

To test whether we could regulate the pH using light under MAS-NMR conditions in a biomolecular sample, we used uniformly ^15^N labeled GB1 as a model protein. GB1 has several titratable residues that show pH-dependent chemical shift perturbations in the pH range 5.5-3.5 ^[2, 26, 27]^ and does not interact with the photoacid (Figure S8). First, ^15^N-GB1 (2 mM) was added to a photoacid buffer containing the trifluoro-alanine/TFANa pH sensor molecules. This solution was transferred to the 3.2 mm MAS rotor. A 1D ^19^F NMR spectrum was recorded to measure pH, followed by a 2D ^1^H-^15^N-HSQC spectrum to observe the protein. Under full-power (100%) light illumination, the solution pH was 3.7, and the HSQC spectra showed well-dispersed GB1 peaks. To perform the pH titration, we reduced the light intensity in a stepwise fashion, recording ^19^F 1D NMR spectra and a 2D ^1^H-^15^N-HSQC spectrum at each illumination level. The ^19^F NMR were monitored to determine when the pH stabilized after each illumination change, a process that took ∼1.5 hours, and then the 2D HSQC spectrum was acquired. This process was repeated at light intensities of 75%, 50%, 25%, 12%, 6%, and 0%. Figure 3A-C shows the change in ^19^F NMR spectra and corresponding change in pH with light intensity. Each intensity change was separated by at least ∼10 hrs, and the pH at a given intensity remained stable throughout the long 2D acquisitions. The HSQC spectra of GB1 acquired at each illumination level and sample pH show the expected titration pattern, with chemical shift perturbations observed for residues T11, K10, D40, E56, G41, among others (Figure 3D-G). These changes are consistent with previous studies of GB1 studied under different pH conditions.^[2, 26, 27]^. These results show that NMR pH titration using light-driven photoacids can be achieved by simply adjusting the light intensity without physically accessing or manipulating the sample, greatly simplifying process and increasing the efficiency of performing such a titration using only single sample. Our photoacid is reversible, so the titration can be reversed by increasing or decreasing light intensity, although the rate of pH stabilization after a change in illumination is faster with decreasing light intensity. The reversible nature is important for returning the sample to the initial condition after titration to rigorously confirm that no degradation or other changes in the sample have occurred during the titration.

**Figure 3:**
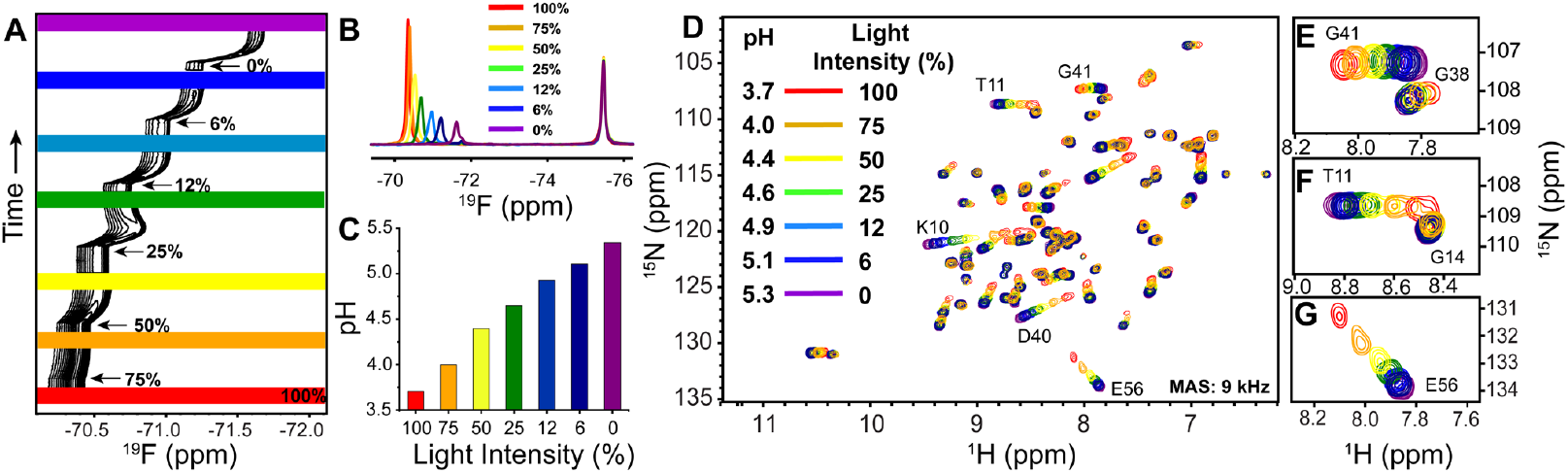
pH titration by varying light intensity: A) Pseudo-2D ^19^F spectra recorded as a function of time under varying light intensities. The colour bars represent the time at which each of the 2D-HSQC spectra in (D) is recorded. B) 1D ^19^F traces showing different pH levels observed at varying light intensities. C) pH measured using the chemical shift differences from the ^19^F spectra shown in (B). D) ^1^H-^15^N HSQC spectra of 2 mM ^15^N-GB1 recorded at different light intensities, showing chemical shift perturbations due to the pH changes. E-G) Zoom-in view of selected residues from (D). All the spectra were acquired with MAS of 9 kHz at sample temperature of 25±5 C at 600 MHz.

The results presented above demonstrate that we can precisely control pH over extended periods, enabling the acquisition of multidimensional solid-state NMR spectra under varying pH conditions. To further explore the applicability of this method, we tested the pKa range of photoacids across different starting pH values. The effective number of protons released depends on the starting pH of the solution. If the starting pH is above the pKa (6.5) the photoacid predominantly exists in its deprotonated (MC) form, minimizing the availability of protons released upon light illumination. This pH range can be extended to near-neutral conditions by decreasing the concentration of buffering components in the solution (Figure S9). We validated this under solution NMR conditions using sub micromolar concentrations of GB1 protein and a pH-sensitive molecule. Upon light illumination, the pH of the solution shifted from 7.0 to 5.2, as confirmed by the HSQC spectra, which showed the expected chemical shift changes corresponding to the pH change (Figure S10). The presence of buffering agents reduces this effect; however, by employing weaker buffering conditions, such as those used in solution NMR, we demonstrated that pH changes from 7.0 to 5.2 could still be achieved. By modifying photoacids, the pH range can be further extended to approach near-neutral conditions. A library of merocyanine-based photoacids is already available in the literature,^[6, 21, 24]^ and adapting them for use under NMR conditions would require straightforward modifications, such as introducing specific functional groups to improve solubility as we have demonstrated here. Furthermore, the precise control of light intensity enables continuous titration and acquisition of a larger number of closely spaced titration points, increasing the accuracy of pKa determination and allowing the extraction of multiple pKa values in complex systems.

In conclusion, we demonstrate that illumination through the top cap of a solid-state MAS rotor can be easily implemented using a 3D-printed adapter and shift of tachometer wavelength, requiring only minimal modification to the MAS NMR probe. Second, we synthesized a photoacid capable of changing the pH upon illumination even under the strong buffering conditions present in NMR samples with high concentrations of biological macromolecules. Third, we show that pH can be controlled by adjusting the light intensity under MAS NMR conditions and different pH values can be stably maintained long enough for recording 2D NMR spectra with a single sample. We believe this method will open new avenues for studying other biological samples with solid-state NMR, as well as solution NMR, and will have broad applications in biological research.

## Supporting Information

Materials and methods, ^1^H NMR and ^13^C NMR and UV-vis spectra of photoacid are included in the supporting information file.

## Acknowledgements

We thank Dr. Songlin Wang, Dr. Paulo Falco Cobra, Dr. Alex Paterson, Boden Vanderloop, Vilius Kurauskas, Ashley Hiett for various facility and sample related help. This work was supported by the National Institute of Health (P41GM136463).

## Notes

Supporting information for this article is given via a link at the end of the document.

### Competing Interest Statement

The authors have declared no competing interest.

