## Supplementary material for "In situ light-driven pH modulation for NMR studies": SI

**Materials and Methods:**

Hydrochloric Acid (HCl), Sodium Hydroxide (NaOH), Methanol (MeOH)-D4, Dichloromethane (DCM), Ethyl acetate (EtOAc), Hexane, Acetonitrile (MeCN), 1,3-propanesultone and 4-Methoxyazobenzene are purchased form Sigma-Aldrich , *p*-methoxyphenylhydrazine hydrochloride and 3-methyl-2-butanone was purchased form Fisher Scientific. Sodium 3-formyl-4-hydroxybenzenesulfonate was purchased from Neta Scientific, $D_2O$ is purchased from Cambridge Isotopes. All the purchased chemicals were used as received without any purification. 980 nm and 365 nm LEDs were purchased from the Mightex® and Multi-Wavelength Fiber Coupled LEDs purchased from Prizmatix Ltd.

**1. Measuring the pKa of the photoacid using UV-vis spectrophotometer:** UV-vis spectra were acquired using a DeNovix DS-11 FX+ UV-Vis spectrophotometer with a DeNovix quartz cuvette (10 mm path length). All experiments were performed in a room lit with Red Fluorescent Tube light (Philips 36W). The sample was prepared by dissolving the photoacid in 20 mL of Milli-Q (MQ) water to a concentration of ~50 µM. The pH of the solution was measured using a Thermo Scientific Orion Star T940 all-in-one titrator equipped with a Hanna Instruments HI1093 extended-length pH electrode with a micro bulb. The pH was adjusted using 100 mM HCl and NaOH. Once a stable pH reading was obtained, 1 mL of the sample was drawn to record the UV spectra. After measuring the UV spectra, the solution was returned for the next titration point. This process was repeated for all the pH points shown in the figure S4. The UV spectra were exported, and absorbance values at 434 nm and 528 nm were plotted in OriginLab for the sigmoidal fit to estimate the pKa.

**2. 3D Printing:** To model the adapter that holds the optical fiber in place, we first disassembled the MAS stator from the Phoenix NMR probe and took all the necessary measurements. Using these measurements, we made a CAD model of the MAS stator in OnShape, a cloud-based CAD platform, and the adapter was modeled to fit on the opening of the stator. The main body of the adapter was printed using a Formlabs Form 3+ printer. The 3D models were downloaded as STL files and uploaded to the Formlabs PreForm software for printing. PreForm software was used to generate the supports and set the orientation. The minimal layer thickness (25 µm) was used for the prints in Durable resin (Formlabs Inc.). The model orientation was optimized to minimize deformation due to the presence of supports.

After printing, excess resin was removed by washing the parts in 99% isopropanol for 15 minutes while still attached to the build plate, and for another 15 minutes after removing them from the build plate and placing them in the wash basket. Tumbling in the basket allows for a more thorough wash. If needed, supports can be removed between steps if the parts are not getting fully washed, allowing IPA to reach areas blocked by the supports. The prints were allowed to dry completely for 60 minutes in a fume hood. We used flush-cut pliers to snap off the supports, and a knife to cut or scrape off any bumps left from the contact points. Sandpaper and files were used to smooth surfaces made rough by the supports. The through-hole for accommodating the optical fiber tended to be partially blocked during printing due to its small diameter, and was expanded out with a hand drill for the optical fiber to fit in smoothly. The holes for the 0-80 nylon screws holding the optical fiber in the adapter and holding the adapter on the stator were manually tapped out as well.

**3. Probe design for the in-situ illumination:** We used a 3.2 mm HFXY probe purchased from Phoenix NMR, which was further modified for *in-situ* illumination purposes. A Prizmatix high-power fiber-coupled four-LED light source with white light, 400 nm, 500 nm, and 700 nm output was purchased from Goldstone Scientific (Southfield, MI). A 980 nm LED light source was purchased from MightX Systems (Pleasanton, CA). Ten-meter-long fiber optics with 400 μm core diameters were purchased from Prizmatix. To illuminate the sample from the top cap, poly(methyl methacrylate) (PMMA) rotor caps for 3.2 mm pencil-style rotors were purchased from Phoenix NMR (Loveland, CO). PMMA was chosen for its high transmittance across all wavelengths tested, from visible to UV ranges.

To hold the optical fiber in place for sample illumination, we designed an adapter that securely holds the optical fiber. A 3D rendering of the adapter design is shown in Figure 1C and Figure S1. The adapter keeps the optical fiber in place and ensures the fiber end is as close as possible to the rotor top cap without touching it. The optical fiber is secured by two nylon screws, and an additional two screws attach the adapter to the stator. The adapter also has holes that function as exhausts for the VT, bearing, and drive gas. Two holes were drilled in the probe cap to accommodate the two optical fibers: one for light illumination and the other for the tachometer wavelength. These two fibers were fed through the upper barrel of the magnet from the top, as shown in Figure S1. For changing the tachometer wavelength, we removed the original optical fiber supplies red tachometer light from the MAS stator and replaced it with the optical fiber connected to the 980 nm LED.

**4. 3.2 mm rotor packing:** PMMA caps (purchased form Phoenix NMR) were inserted into the rotor and glued around the edges of the caps. Any excess glue was removed by wiping it along the outsides, and the caps were allowed to cure for 30 minutes. Sample packing involved transferring 20 μL of solution using a sharp pipette tip, after which the rotor was closed with a silicon rubber disc, followed by inserting the drive tip. For tach marking, the rotor was initially marked with silver on all sides of the top cap, and after a 5-minute drying period, the rotor cap was painted with a black Sharpie. This was followed by another 5 minutes of drying. This marking process was repeated three times, and in the final stage, half of the rotor was marked black and the other half with silver.

**5. MOAB-Sample preparation:** A solution of MOAB in deuterated methanol at a concentration of 10 mg/mL was packed in a standard wall 3.2 mm pencil-style rotor with PMMA caps as described above. For illumination experiments of MOAB, the rotor was spun at 10 kHz. 365 nm UV light was used to trigger the conformational change of the MOAB double bond.

**6. Photoacid sample preparation:** 4 mg of photoacid was dissolved in 0.2 mL of buffer containing 10 mM sodium trifluoroacetate. The pH was adjusted to 5.3 using 100 mM Sodium hydroxide. All the preparation were done in a room lit with Red Fluorescent Tube light. 20 μL of solution was transferred to a 3.2 mm rotor with PMMA caps glued on one side and other side is closed with the silicon rubber disc and drive tip.

**7. GB1 Sample Preparation:**$^{15}$N-GB1 was first exchanged from the storage buffer with a buffer containing 10 mM sodium trifluoroacetate, 10 mM trifluoro alanine, and 50 mM sodium chloride, adjusted to pH 5.3 using Amicon centrifugal concentrators. After the buffer exchange, the sample was concentrated to obtain a 4 mM GB1 solution. A 40 mM photoacid solution was prepared by dissolving photoacid powder in a 10 mM sodium trifluoroacetate solution, with the pH adjusted using 100 mM NaOH. The NMR sample was prepared by adding 20 μL of the photoacid solution to 20 μL of the GB1 solution. Finally, 20 μL of the final solution was transferred to the 3.2 mm rotor and the rotor was sealed as mentioned previously.

**8. NMR Spectroscopy:** All magic-angle spinning NMR experiments were performed on a 600 MHz instrument equipped with an AVANCE III HD console. Solution NMR experiments were performed on 600 MHz and 500 MHz instruments, also equipped with AVANCE III HD consoles.

**8.1. Solution 1D $^{1}$H and 1D $^{13}$C NMR:** Solution 1D experiments were conducted on a 500 MHz instrument equipped with a 5 mm C13-optimized TXO cryoprobe. $^{1}$H NMR spectra were recorded using a one-pulse experiment, while the $^{13}$C spectra were recorded with inverse-gated $^{1}$H decoupling. $^{1}$H spectra were acquired with a recycle delay of 5 seconds and 16 scans. $^{13}$C spectra were obtained using a 30° flip angle pulse with $^{1}$H decoupling during acquisition, with 512 scans.

**8.2 Solution $^{19}$F NMR:** 1D $^{19}$F experiments were performed using a 5 mm QCI-F cryoprobe ($^{1}$H/$^{19}$F/$^{13}$C/$^{15}$N) on a 600 MHz instrument equipped with an AVANCE III HD console. $^{19}$F was observed with inverse-gated $^{1}$H decoupling. $D_2O$ was used for locking. The spectra width was 29.5 ppm, offset -70 ppm, with 16 scans, a pulse width of 12.2 μs, a recycle delay of 1.5 seconds, an acquisition time of 1 second, and $^{1}$H decoupling using WALTZ64.

**8.3 Magic-angle spinning 1D $^{1}$H NMR:** Solid-state 1D experiments were performed at 10 kHz MAS with a sample temperature of 25°C, using a Hahn echo with pre-saturation of water during the recycle delay. Other parameters: 32 scans, 2-second recycle delay, 1-second acquisition time, spectral width of 16 ppm, a 5 μs $^{1}$H pulse, an echo of four rotor periods, and pre-saturation with a pulse power of 280 Hz for 2 seconds.

**8.4 Magic-angle spinning $^{19}$F NMR:** 1D $^{19}$F spectra were acquired using inverse-gated $^{1}$H decoupling with a Phoenix Probe configured in HFCN, according to the manufacturer's recommendations. Spectra were acquired with the following parameters: 32 scans, 1-second acquisition time, $^{19}$F offset of -70 ppm, spectral width of 80 ppm, a 5 μs $^{19}$F pulse, 2-second recycle delay, and continuous wave $^{1}$H decoupling of 1.5 kHz applied during acquisition.

**8.5 2D $^{1}$H-$^{15}$N Heteronuclear Single-Quantum Coherence (HSQC):** All 2D experiments were performed using the Bruker pulse program "*hsqcphpr*" The sample was spun at 9 kHz with a temperature of 25°C. Water suppression was achieved using a pre-saturation pulse of 280 Hz during the recycle delay of 1.5 sec. Polarization transfer obtained with INEPT delay of 2.77 ms ($^{1}J_{H-N}$ = 90 Hz). Other parameters: 64 scans, td 1998x200, acquisition time of 100 ms ($t_2$) and 22 ms ($t_1$), spectral width of 16.6 ppm ($^{1}$H) and 74 ppm ($^{15}$N), $^{1}$H pulse width of 4.8 μs, and $^{15}$N pulse width of 6.4 μs. All spectra were processed using NMRPipe and analyzed using NMRFAM-Sparky.

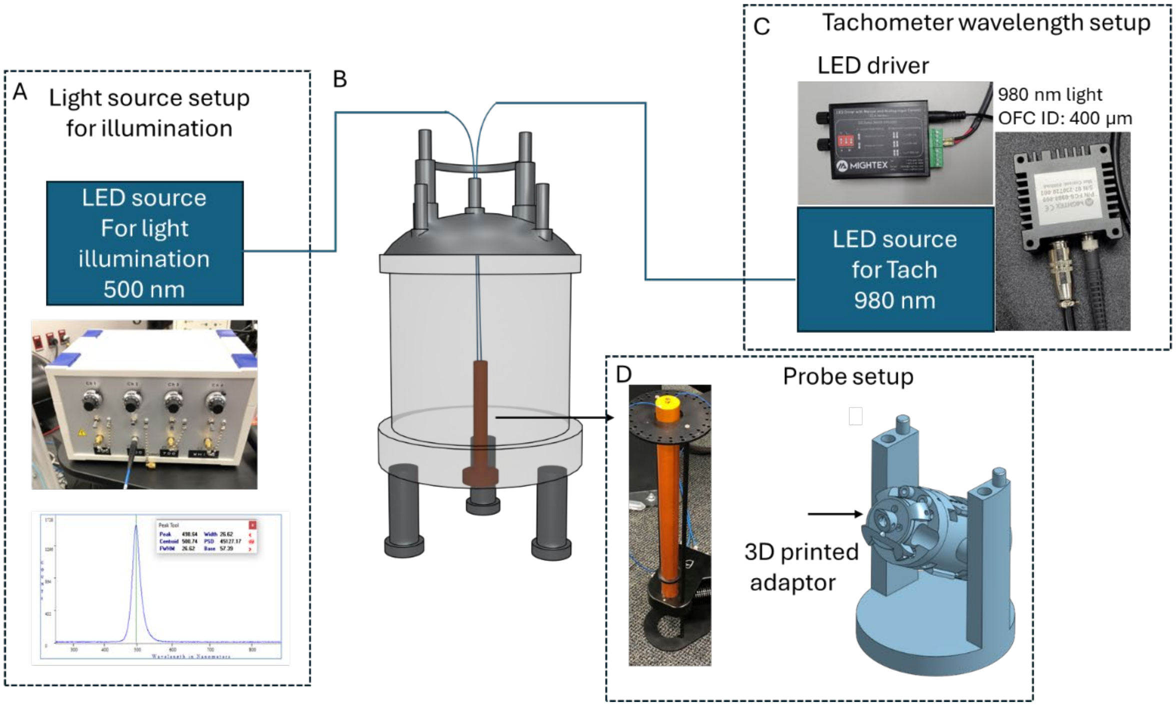


**Figure S1: Illustration of the photoprobe setup:** A) Light source for sample illumination. The inset shows the LED device, 500 nm emission spectrum (bottom), as provided in the manufacturer's manual. B) Schematic representation of the NMR magnet showing the fiber optics setup. C) Picture showing the LED source and LED driver used for the tachometer. D) Picture of the Phoenix probe showing the optical fibers emerging from the probe, with two holes drilled into the cap. A 3D design shows the adapter and screw holes for securing the adapter onto the MAS stator.

**9. Synthesis of photoacid.** The photoacid was synthesized by following the literature procedure.[1-3] Briefly In a 250 mL round-bottom flask, p-methoxyphenylhydrazine hydrochloride (5 g, 20.3 mmol) and 3-methyl-2-butanone (10 g, 21.6 mmol) were dissolved in glacial acetic acid (30 mL). This solution was then heated under reflux for 16 hours. The solution was left to cool, allowing the products to separate, and the solvent was decanted. A brown residue was dissolved in dichloromethane (DCM) and washed with saturated sodium sulfate. The final solution was purified by flash chromatography (silica, hexane: ethyl acetate (EtOAc) 2:1). Fractions containing a yellow residue were collected.

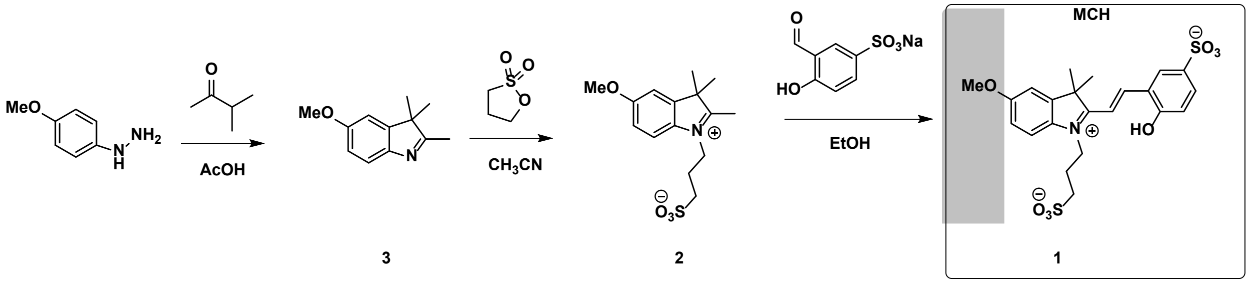


**Scheme S1. Synthetic route for the synthesis of photoacid 1.**

The obtained yellow residue was mixed in another 250 mL round-bottom flask with 1,3-propanesultone. These were dissolved in acetonitrile (MeCN) and heated under reflux for 16 hours. The resulting brown solution was added dropwise to EtOAc under stirring. After allowing it to rest, a beige precipitate formed, which was separated by vacuum filtration and washed with EtOAc. The precipitate was recrystallized from hot MeCN and left overnight in a -20°C freezer. After decanting, purple crystals were obtained.

In a 100 mL round-bottom flask, the purple crystals (1.2 g, 20.1 mmol) were dissolved in ethanol (~20 mg/mL). Sodium 3-formyl-4-hydroxybenzenesulfonate was purchased commercially and (2 equivalents) was added to the ethanol solution, and the mixture was heated to reflux for 16 hours. The solution was then transferred into two 50 mL centrifuge tubes for centrifugation to separate the ethanol from the precipitate. The orange precipitate was washed with ethanol. The product was dissolved in water and lyophilized. The resulting product was characterized by $^{1}H$ and $^{13}C$ NMR.

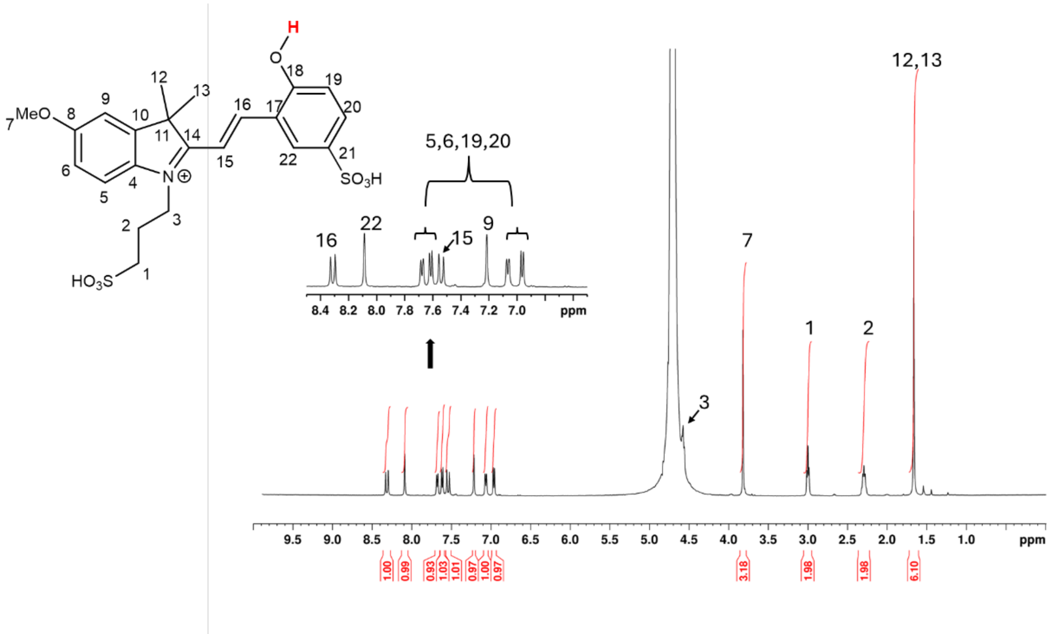


**Figure S2: Characterization of photoacid using $^{1}$H NMR:** $^{1}$H NMR spectra of the photoacid **1** showing the integrated area for each proton. The integrated areas match with the expected structure. Conditions used: 20 mM in $D_2O$ and pH is adjusted to ~2.0 using 1 M HCl. The spectrum was recorded under low pH condition to avoid multiple conformations of the photoacid. Under these conditions only the protonated MCH form is present in solution.

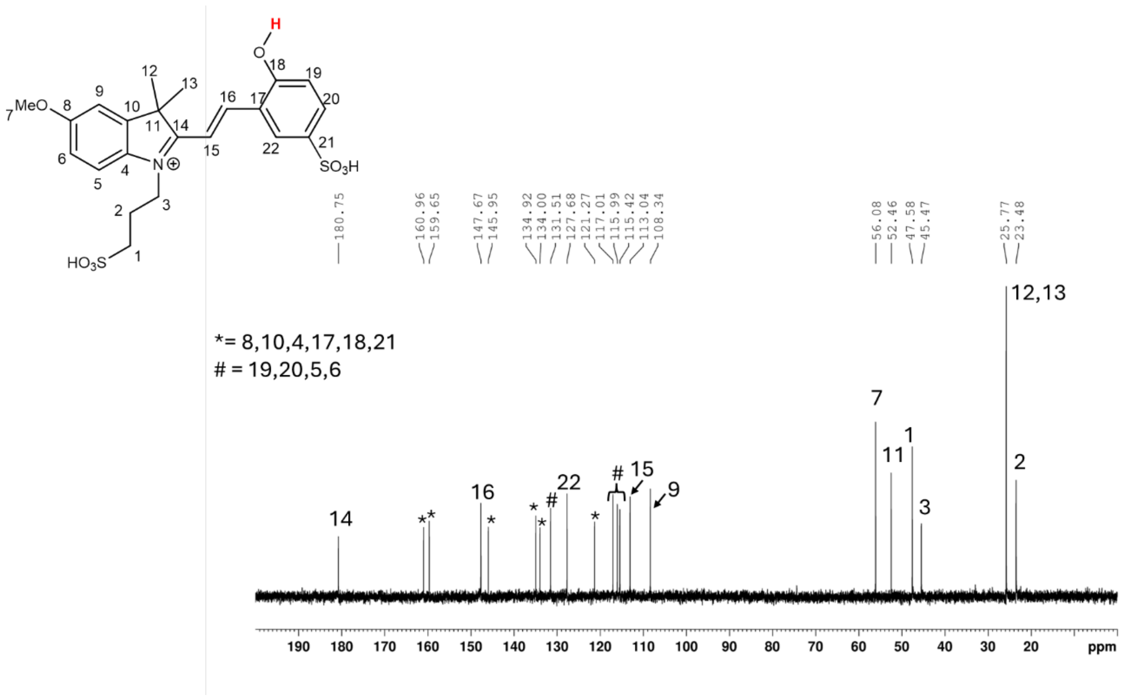


**Figure S3: Characterization of photoacid using $^{13}$C NMR:** $^{13}$C NMR spectra of the photoacid **1**. Conditions used: 20 mM in $D_2O$ with the pH adjusted to ~2.0 using 1M HCl.

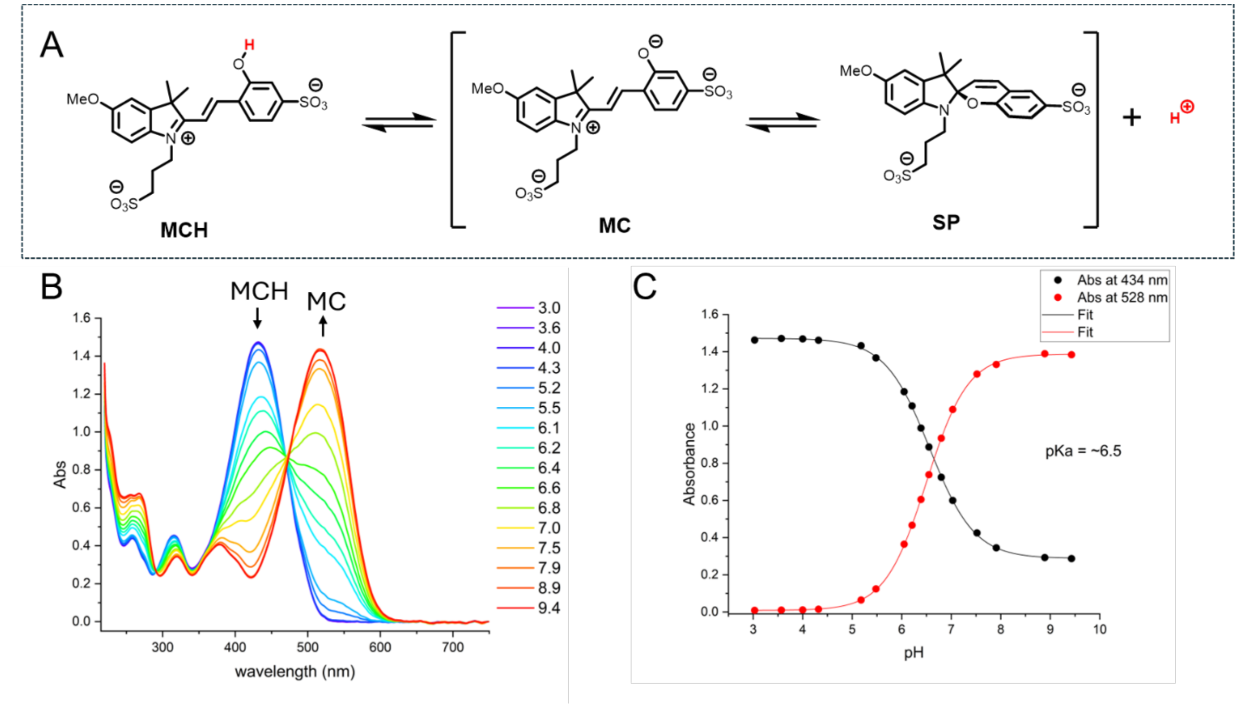


**Figure S4: Measurement of pKa of photoacid 1:** A) Chemical structure of the photoacid equilibrium in the dark between the protonated merocyanine form (MCH), the deprotonated form (MC) and respective ring-closed product of MC – the spiropyran form (SP), to determine the pKa under dark conditions. B) UV-Vis absorption spectra of the photoacid measured in the dark at different pH values as shown in the figure. C) Absorption of MCH (at 434 nm) and MC (at 528 nm) plotted against pH. The sigmoidal fit shows a pKa of 6.5 for the molecules under dark conditions. Concentration ~50 µM. The pKa measured here represents pseudo pKa values due to the equilibrium between the MC and SP forms but are useful in estimating the effective concentration of the active MCH form.

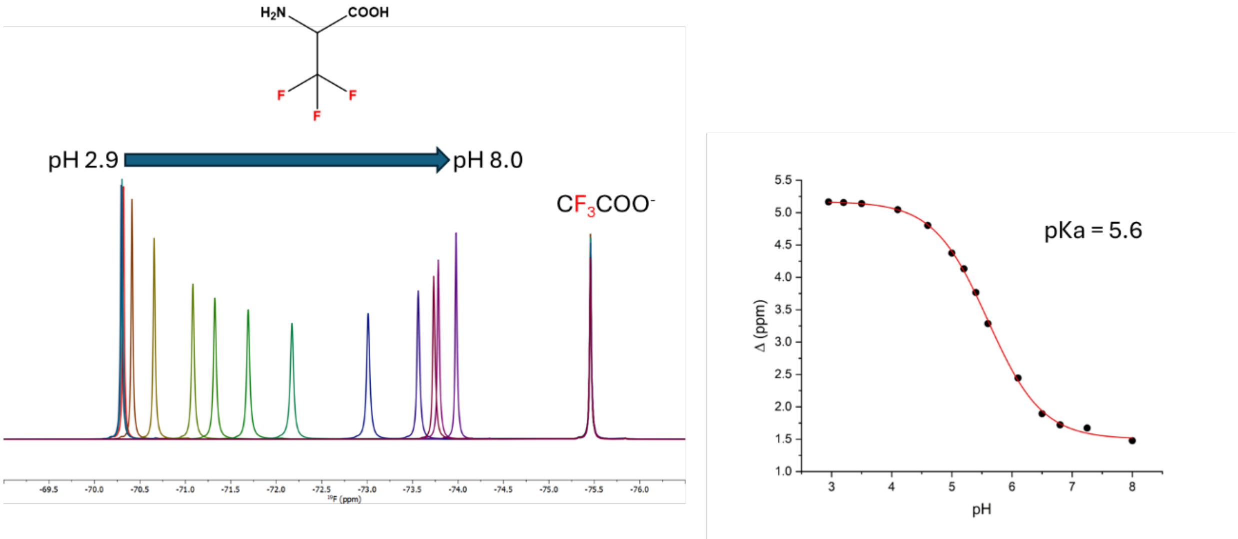


**Figure S5: pKa measurement of trifluoro-alanine:** [19]F 1D-NMR of *trifluoroalanine* at different pH values. *Sodium trifluoroacetate* was used as an internal chemical shift standard. The difference in chemical shift was measured and plotted against pH in the figure on the right. The sigmoidal fit shows the pKa of *trifluoroalanine*. Using this standard curve, the pH of the solution can be obtained by measuring the chemical shift difference between *trifluoroalanine* and sodium *trifluoroacetate*.

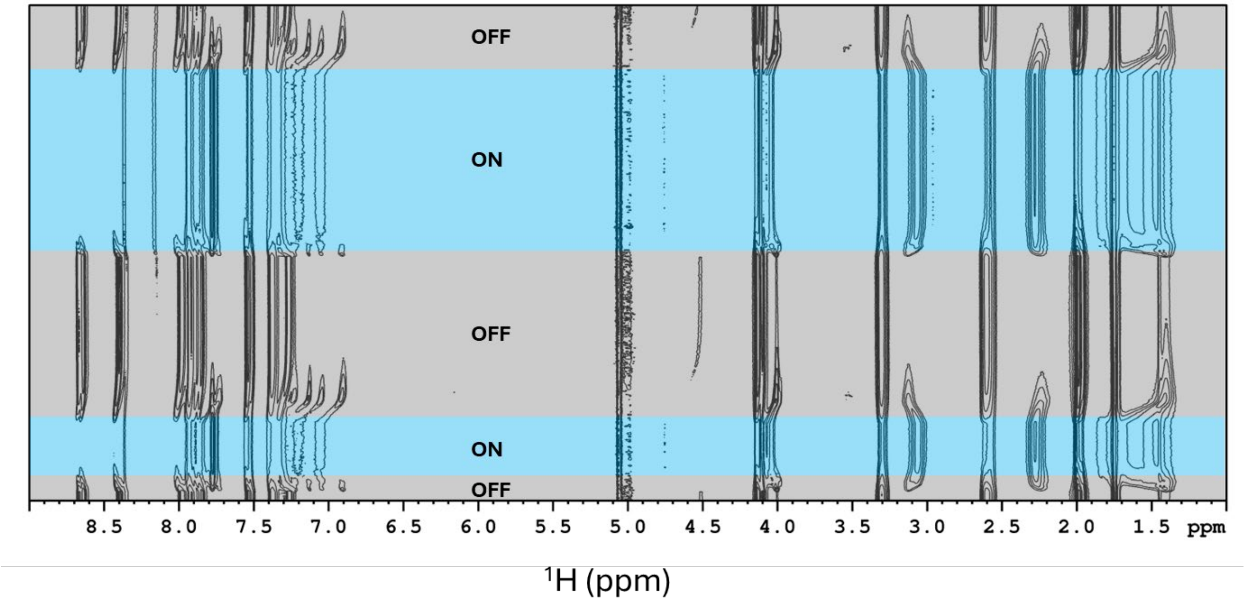


**Figure S6: $^1$H NMR of photoacid conversion with 500nm light:** Pseudo-2D $^1$H spectra recorded as a function of time with the light ON and OFF as indicated. The $^1$H spectra show complete conversion of the photoacid SP conformation upon illumination, which reverts back to the MCH form when the light is turned OFF.

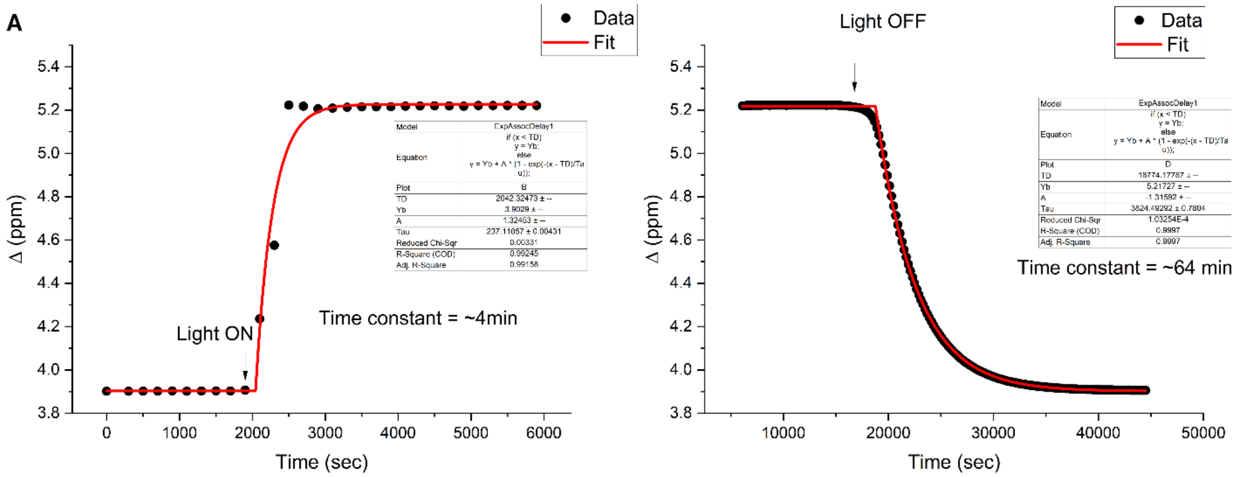


**Figure S7:** A) Plot showing the changes in the chemical shift difference between trifluoro-alanine and TFANa after light illumination as a function of time. The data is fitted to an exponential function with a delay component to extract the time constant for the chemical shift changes upon illumination. All parameters and the equation used for the fit are shown in the box. The plot indicates that the change upon illumination has a time constant of approximately 4 minutes. B) Time constant measured to reach equilibrium under dark conditions upon switching off the light (~64 minutes). All fits were performed using OriginLab software with default settings, and the parameters are shown in the box. These data are also presented in Figure 2C of the main text.

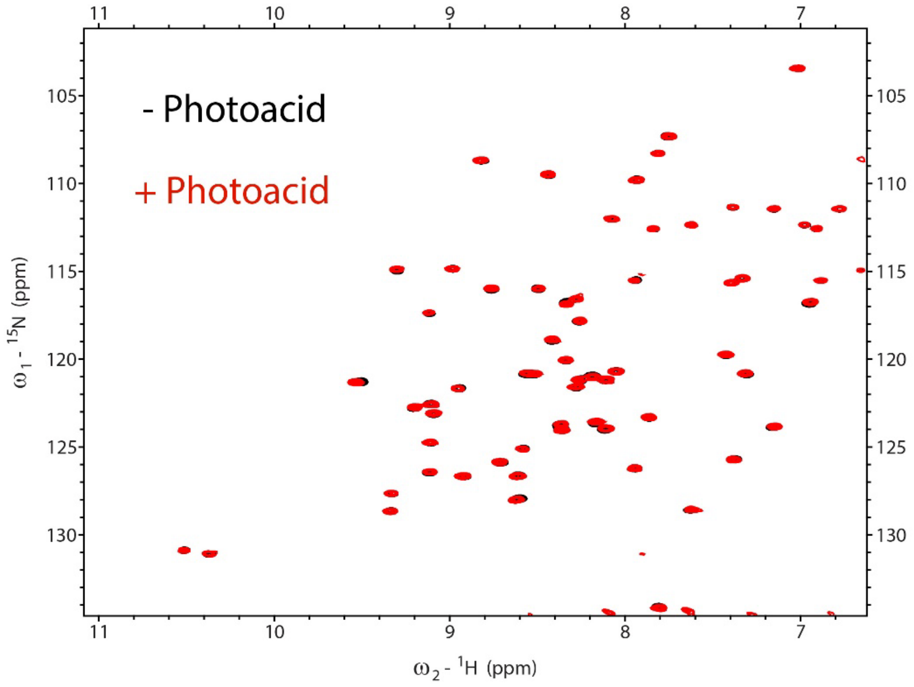


**Figure S8:** $^1$H-$^{15}$N HSQC spectra of $^{15}$N-GB1 in 50 mM NaCl recorded with (red) and without (black) photoacid (20 mM) showing that the photoacid is not interacting with GB1. These experiments were performed using a solution NMR spectrometer with 0.2 mM $^{15}$N-GB1, 20 mM Photoacid and 50 mM NaCl.

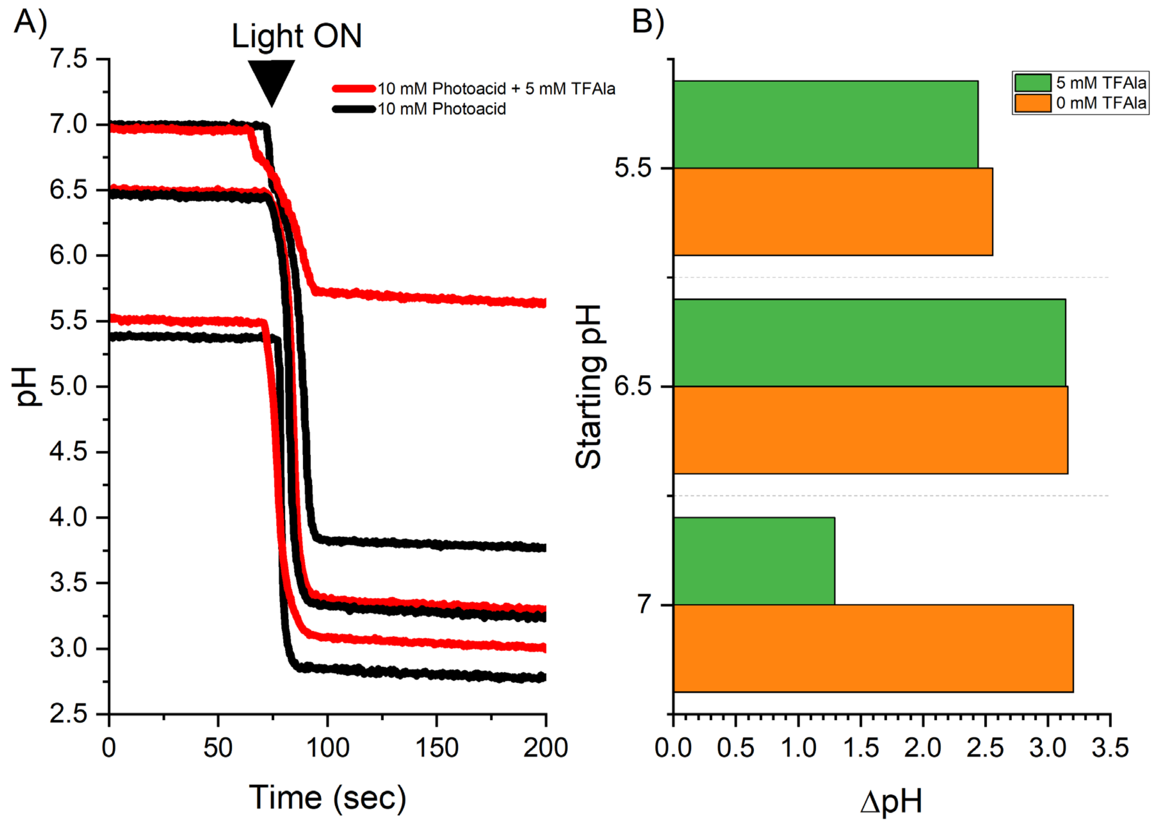


**Figure S9:** Effect of initial pH and the presence of buffering molecules on the photoacid working range. (A) Change in pH upon light illumination as a function of initial pH under varying starting conditions, with and without 5 mM TFAla. (B) Plot showing the effective pH change observed across different initial pH values in the presence and absence of 5 mM TFAla. These results indicate that the working range of the photoacid is strongly dependent on both the presence of a buffer and the initial pH. The experiments were performed using white flashlight illumination, and pH measurements were obtained using a custom-built setup designed to monitor pH changes over time.

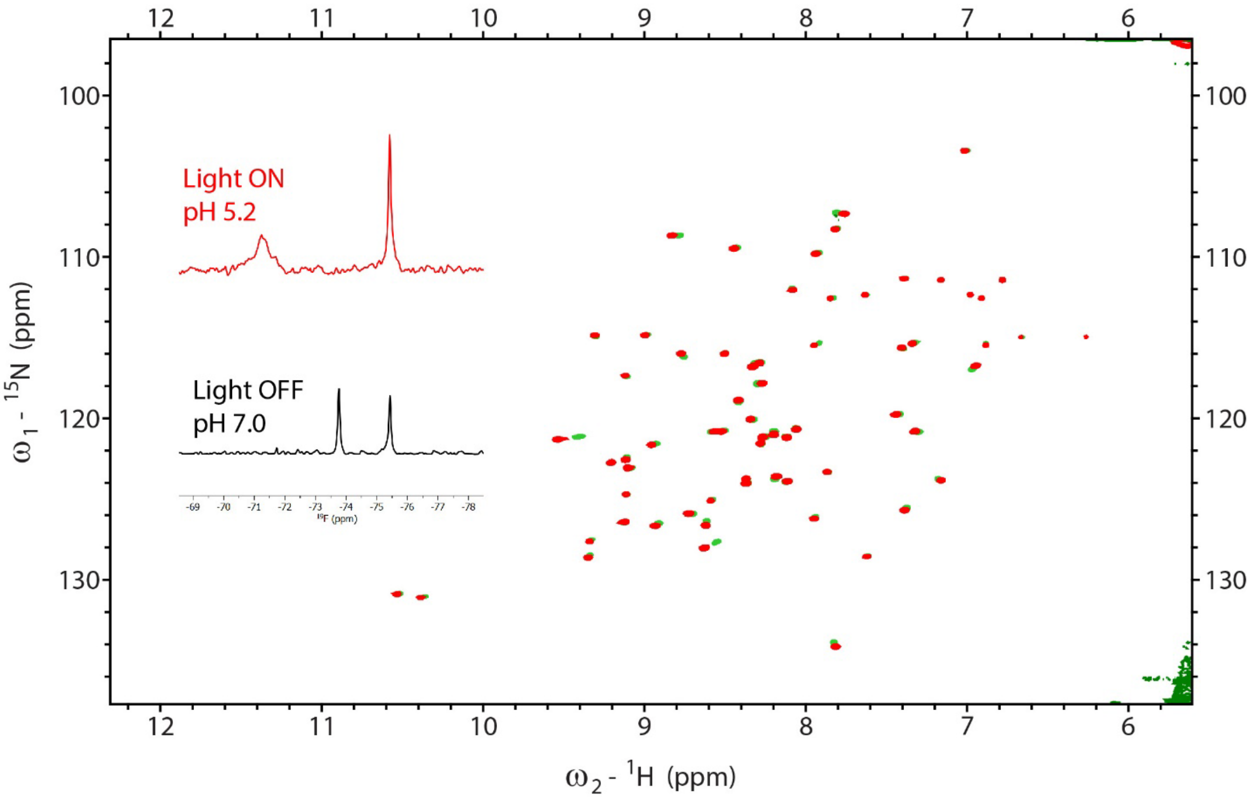


**Figure S10:** $^{1}H$-$^{15}N$ HSQC spectra recorded with the photoacid in dark conditions (black) and under illumination with 500 nm light (red), highlighting chemical shift perturbations caused by pH changes. Experimental conditions included 100 μM $^{15}N$-GB1, 20 mM photoacid, 0.1 mM TFAla, and 0.1 mM TFANa, with a starting pH of 7.0. These experiments were performed using solution NMR spectroscopy with a 3 mm solution NMR tube. Light illumination was achieved by directly inserting an optical fiber into the NMR tube. The tip of the optical fiber was sanded to enhance sample illumination throughout the solution. Inset showing the $^{19}F$ 1D traces used to measure the pH changes upon illumination with light. All spectra were recorded at a sample temperature of 25°C using a 600 MHz spectrometer equipped with a Bruker Avance III HD console and a QCI-F (1H/19F/13C/15N) cryoprobe with Z-axis gradients.

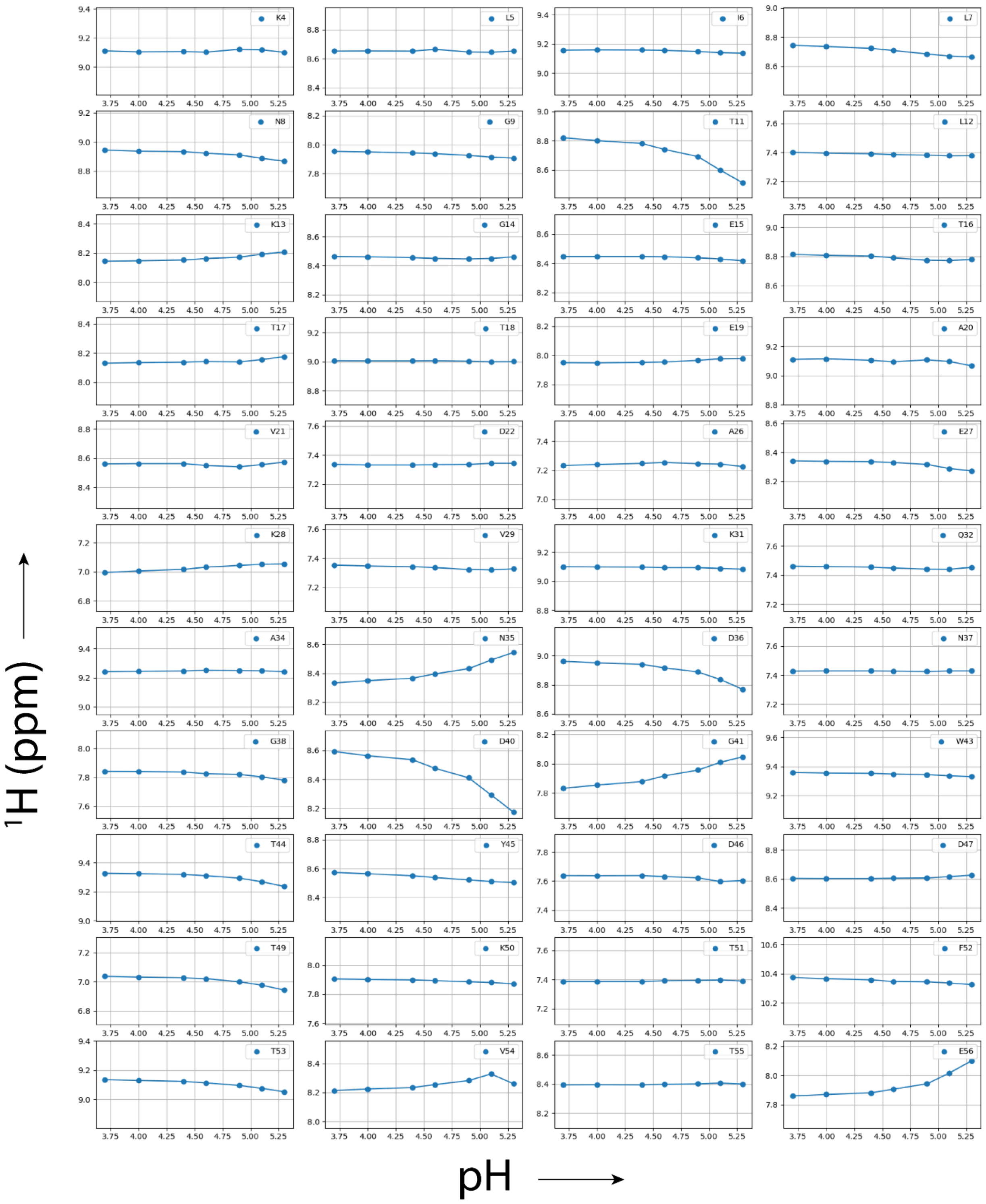


**Figure S11:** $^{1}$H Chemical shifts changes for each residue as function of pH measured form the HSQC spectra presented in figure 3.

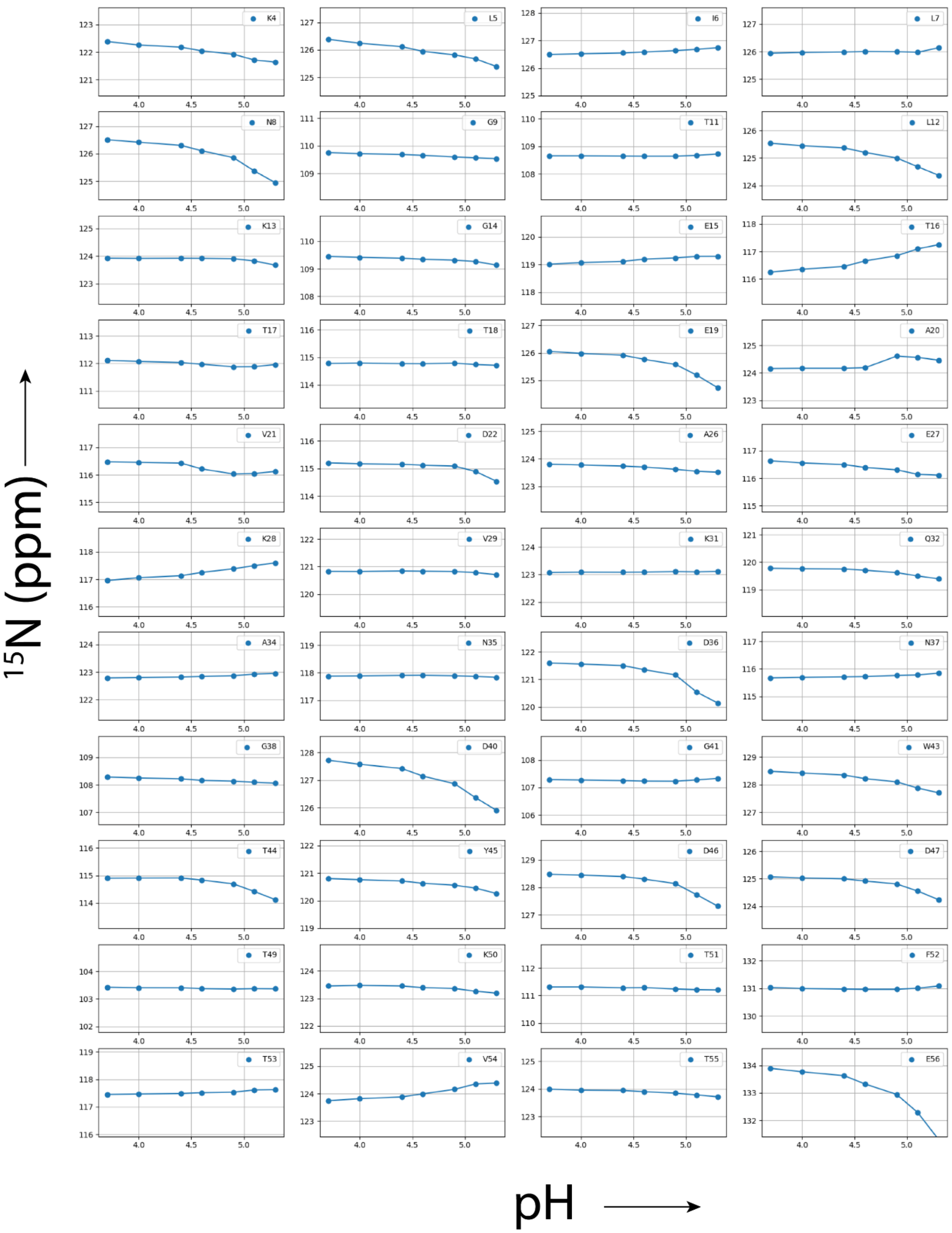


**Figure S12:** $^{15}$N Chemical shifts changes for each residue as function of pH measured form the HSQC spectra presented in figure 3.
